# Spectrin forms a periodic cytoskeleton within the epidermis to preserve axonal integrity

**DOI:** 10.1101/2022.09.07.507048

**Authors:** Sean Coakley, Igor Bonacossa-Pereira, Massimo A. Hilliard

## Abstract

Spectrins are highly conserved molecules that form a distinct membrane associated periodic scaffold within axons thought to provide mechanical resilience. In *C. elegans*, UNC-70/ß-Spectrin also functions outside the nervous system, within the epidermis, to maintain the integrity of sensory neurons. The precise molecular organization and cellular mechanisms that mediate this protection are unknown. Here, using 3D-structured illumination microscopy, we show that within the epidermis spectrins form a crescent-shaped scaffold with a periodicity of ∼ 200 nm that embraces adjacent axons. This epidermal spectrin scaffold is induced by developing axons and reformed during axonal regeneration, and creates a “molecular imprint” of the adjacent nervous system. Disruption of this epidermal scaffold causes widespread axonal damage in sensory and motor neurons. These findings reveal the existence of a distinct and periodic spectrin framework within the epidermis that is molded by the developing nervous system and is necessary to protect it from mechanical damage.

**Teaser:** Spectrin forms a periodic scaffold in the epidermis that protects axons from damage.

## MAIN TEXT

The ubiquitous axonal membrane-associated periodic scaffold (MPS) has been proposed to provide axons with an intrinsic ability to withstand strain. In the last decade, since first being revealed by super-resolution microscopy, its unique structure of actin rings interspaced with α- and ß-Spectrin tetramers with a periodicity of ∼200 nm has been observed within axons of every neuronal subtype and species tested ^1,2^. In the nematode *C. elegans*, mutations in *unc-70/ß-Spectrin* cause sensory and motor axons to spontaneously break due to the strain of body movement ^3,7-9^. In addition, spectrins have been shown to provide tension to axons ^3^ and contribute to resisting mechanical stress ^4,5^. Recent work has hinted that non-neuronal spectrin may play a larger role than previously recognized to protect neurons from damage. We recently demonstrated that ß-Spectrin functions in the epidermis to protect mechanosensory neurons of *C. elegans* from damage ^7^. However, the organization of spectrin within this tissue and how it protects from axonal damage is unclear. Here, we investigate the nanoscale architecture of epidermal spectrin and interrogate its role in maintaining axonal integrity.

Endogenous *ß-Spectrin* (GFP::UNC-70) and *a-Spectrin* (SPC-1::GFP) are expressed throughout *C. elegans* and enriched in axons (Fig. 1a-c)^10,11^. To visualize the localization of endogenous UNC-70/ß-Spectrin selectively within the epidermis, we adopted a split fluorophore strategy using split GFP separated between the 10^th^ and 11^th^ ß-strands to generate two fragments: spGFP_1-10_ and spGFP_11_. Using CRISPR-Cas9 we fused three tandem copies of spGFP_11_ to the endogenous N-terminus of UNC-70/ß-Spectrin. We then expressed spGFP_1-10_ selectively within the epidermis (SKIN::spGFP_1-10_ = *Pdpy-7::spGFP*_*1-*10_) and observed reconstitution of full-length epidermal GFP_x3_::UNC-70. Strikingly, this labeled the epidermal membrane surrounding axons, revealing an “imprint” of the nervous system within the epidermis (Fig. 1d). Using a similar strategy, we visualized the localization of endogenous SPC-1/α-Spectrin using a strain in which seven tandem copies of spGFP_11_ were fused to the C-terminus of SPC-1/α-Spectrin ^11^. As observed with UNC-70/ß-Spectrin, epidermally expressed spGFP_1-10_ (SKIN::spGFP_1-10_ = *Pdpy-7::spGFP*_*1-*10_) resulted in the reconstitution of full-length epidermal SPC-1::GFP_x7_, clearly labeling the same membrane regions of the epidermis in contact with adjacent axons. An imprint of the nervous system was also labeled by epidermal SPC-1::GFP_x7_ and confirmed by visualizing epidermal SPC-1::GFP_x7_ together with a red fluorescent marker of either GABAergic motorneurons or mechanosensory neurons (Fig. 1e-i).

**Fig. 1.**
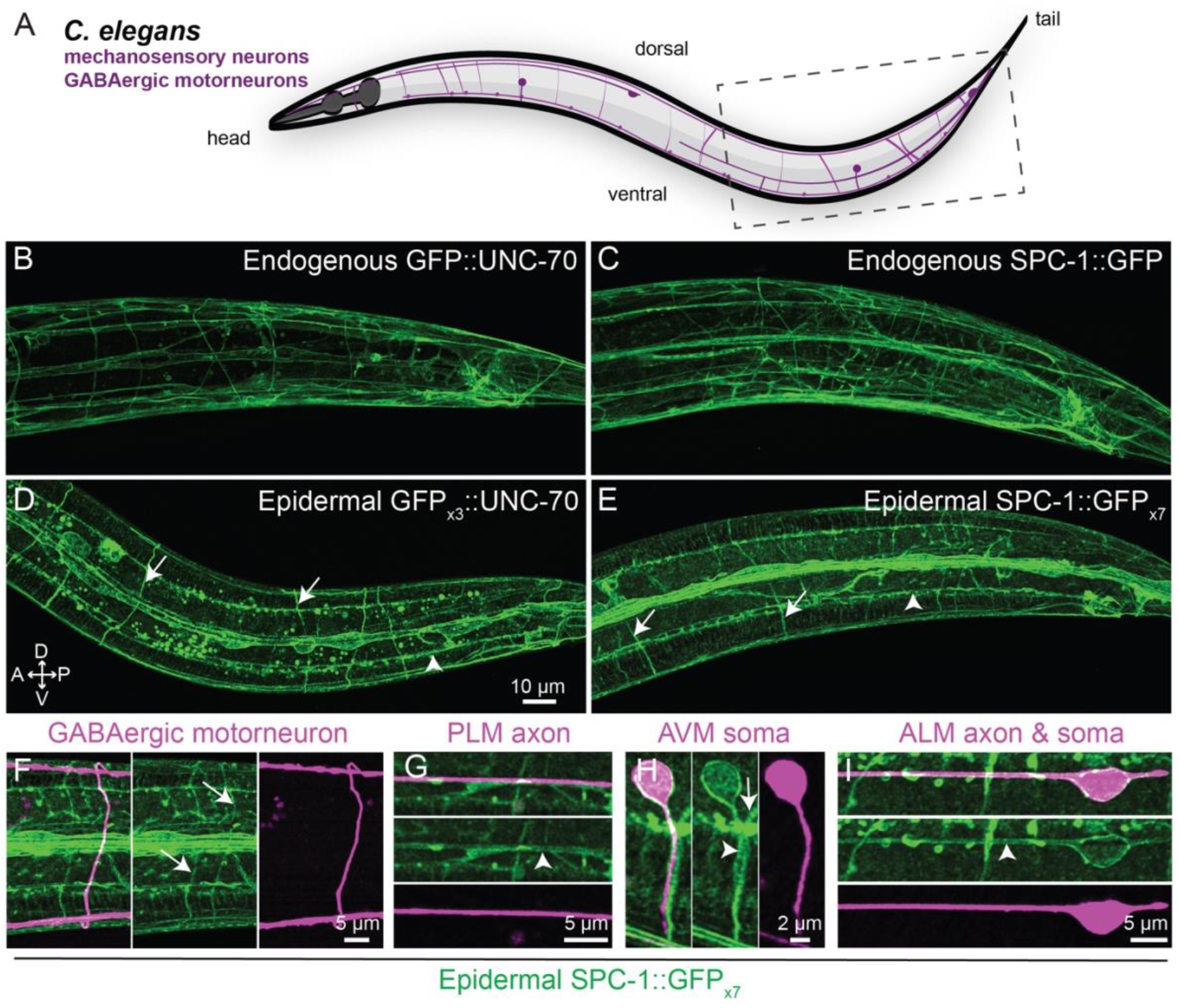
The epidermal spectrin cytoskeleton reveals an imprint of adjacent axons. (A) Schematic of *C. elegans* with mechanosensory and GABAergic motorneurons depicted. (B-E) Lateral view of the tail of a wild-type animal (boxed region in A) at the L4 stage showing the localization of (B) endogenous GFP::UNC-70, (C) endogenous SPC-1::GFP, (D) reconstituted epidermal GFP_x3_::UNC-70 (spGFP_11×3_::UNC-70 + SKIN::spGFP_1-10_), (E) reconstituted epidermal SPC-1::GFP_x7_ (SPC-1::spGFP_11×7_ + SKIN::spGFP_1-10_). (F-I) Localization of reconstituted epidermal SPC-1::GFP_x7_ and a red fluorescent marker labeling (F) GABAergic motorneurons, (G) the PLM axon, (H) the AVM soma and axon, and (I) the ALM soma and axon. In (D-I) arrows mark the outline of motorneuron commissures, and arrowheads mark the outline of the mechanosensory neurons within the epidermis.

To determine whether this epidermal spectrin structure developed before or after axonal extension, we examined its appearance in relation to the GABAergic motorneurons at different stages of development. In the first larval stage after hatching (L1), epidermal SPC-1::GFP_x7_ was not present on GABAergic commissures (Fig. 2a), whereas it appeared by the third larval stage (L3) and clearly labeled commissural axons and motorneuron cell bodies (Fig. 2b-c). To further test the notion that this epidermal spectrin localization was induced by the underlying neurons, we performed axotomies on the PLM mechanosensory neuron and visualized the localization of epidermal SPC-1::GFP_x7_ on the nascent regrowing axon. 48 h after injury, epidermal SPC-1::GFP_x7_ localized to regrowing proximal axons adjacent to the epidermis (Fig S1). Moreover, the degenerating distal axonal fragment, now separated from the cell body, had begun to lose its epidermal SPC-1::GFP_x7_ imprint (Fig S1). Taken together, these data indicate that it is the underlying axon that induces the formation of a spectrin imprint in the adjacent epidermis.

**Fig. 2.**
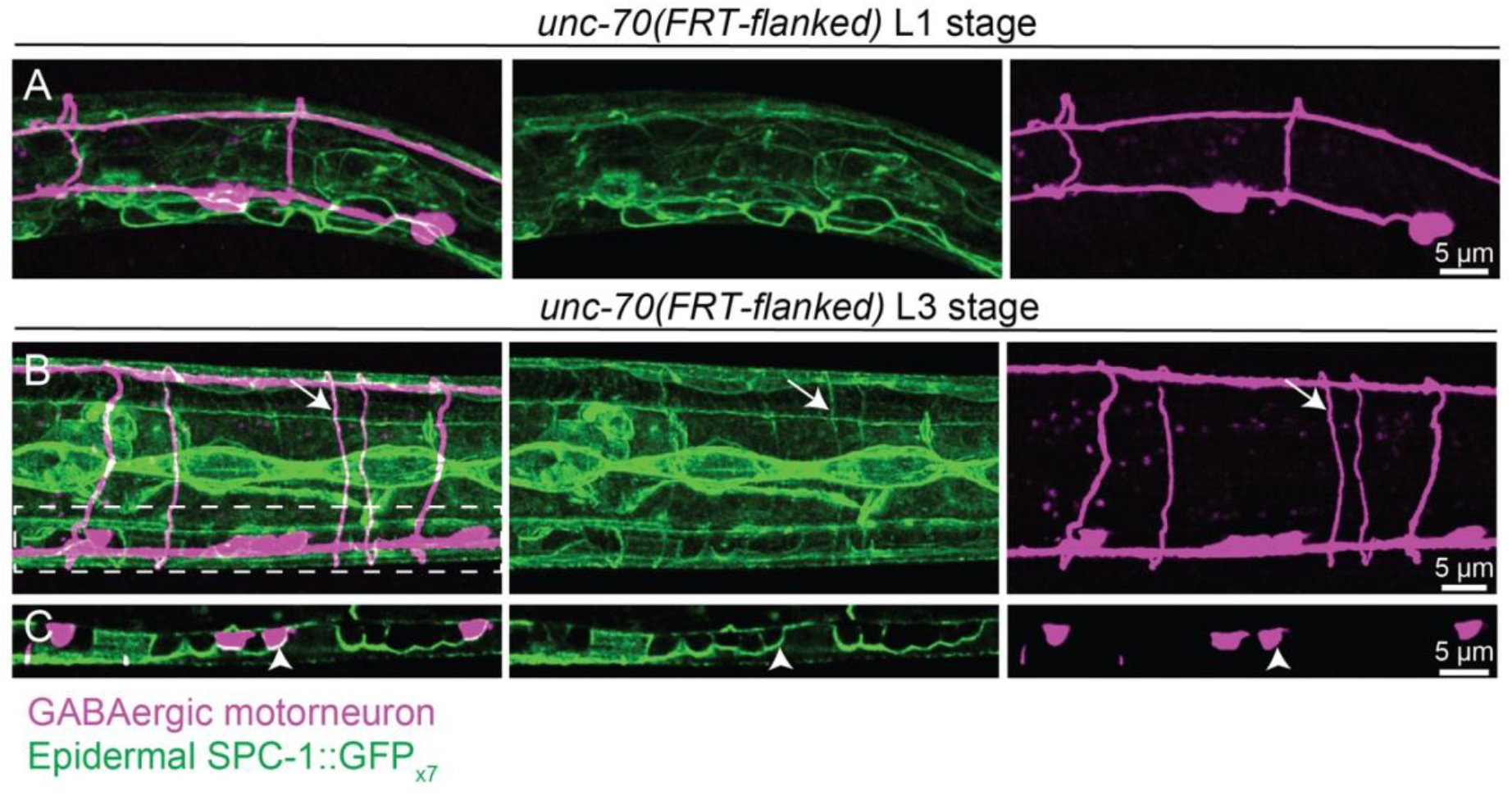
Developing neurons induce an epidermal Spectrin imprint of the nervous system. (A, B) Localization of reconstituted epidermal SPC-1::GFP_x7_ (SPC-1::spGFP_11×7_ + SKIN::spGFP_1-10_) and a red fluorescent marker labeling GABAergic motorneurons at the first larval stage (L1) (A), and at the third larval stage (L3) (B). Arrows indicate the epidermal SPC-1::spGFP_x7_ localization corresponding to a GABAergic motorneuron commissure. (C) Single slice through boxed area shown in B showing the ventral nerve cord and reconstituted epidermal SPC-1::spGFP_x7_ surrounding GABAergic motorneuron cell bodies (arrowheads).

Importantly, the localization of SPC-1::GFP_x7_ surrounding these axons appeared discontinuous and periodic (Fig S2a-c). To confirm this observation and determine the nanoscale organization of this epidermal spectrin network, we visualized epidermal GFP_x3_::UNC-70 and SPC-1::GFP_x7_ using 3D-structured illumination microscopy (3D-SIM). In regions of the epidermis where GFP_x3_::UNC-70 and SPC-1::GFP_x7_ localized to the membrane adjacent to axons, we observed a periodic localization of GFP_x3_::UNC-70 and SPC-1::GFP_x7_ reminiscent of the axonal MPS (Fig. 3a,c, e, & g). This localization was seen surrounding the axons of the ALM and PLM mechanosensory neurons as well as the GABAergic motorneuron commissures, with a period of ∼200 nm (Fig. 3a-h & k, l). Unlike the rings seen within the axonal MPS, the epidermal SPC-1::GFP_x7_ localization presented a crescent shape surrounding one half of the axonal shaft (Fig. 3k).

**Fig. 3.**
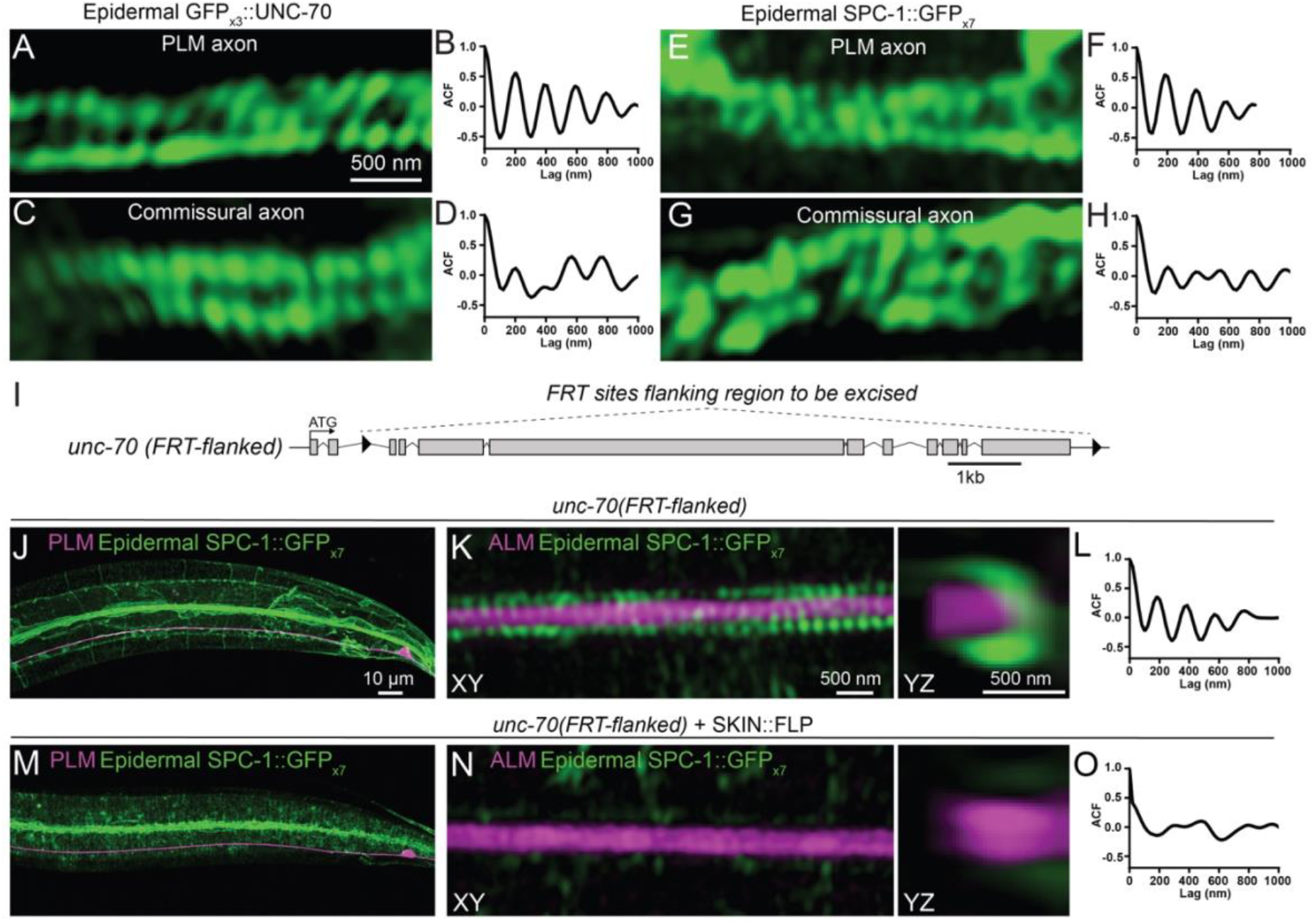
Epidermal spectrins form a periodic membrane-associated scaffold surrounding axons. (A) 3D-SIM image showing localization of reconstituted epidermal GFP_x3_::UNC-70 (spGFP_11×3_::UNC-70 + SKIN::spGFP_1-10_) in a region surrounding the PLM axon. (B) Autocorrelation function of the periodic region shown in A. (C) 3D-SIM image showing the localization of reconstituted epidermal GFP_x3_::UNC-70 in a region surrounding a commissural axon. (D) Autocorrelation function of the periodic region shown in C. (E) 3D-SIM image showing localization of reconstituted epidermal SPC-1::GFP_x7_ (SPC-1::spGFP_11×7_ + SKIN::spGFP_1-10_) in a region surrounding the PLM axon. (F) Autocorrelation function of the periodic region shown in E. (G) 3D-SIM image showing localization of reconstituted epidermal SPC-1::GFP_x7_ specifically in the epidermis in a region surrounding a commissural axon. (H) Autocorrelation function of the periodic region shown in G. (I) A schematic depicting a conditional *unc-70/ß-Spectrin* allele containing flippase recognition target (FRT) sites for tissue-specific flippase-dependent excision. (J) Lateral view of the tail of an *unc-70(FRT-flanked)* animal at the L4 stage showing the localization of reconstituted epidermal SPC-1::GFP_x7_ (SPC-1::spGFP_11×7_ + SKIN::spGFP_1-10_) with a red fluorescent marker labeling the PLM mechanosensory neuron. (K) 3D-SIM image showing localization of reconstituted epidermal SPC-1::GFP_x7_ (SPC-1::spGFP_11×7_ + SKIN::spGFP_1-10_) in a region surrounding the axon of the mechanosensory neuron ALM in an *unc-70(FRT-flanked)* animal at the L4 stage. Left panel is a lateral view, right panel an orthogonal perspective of the projected z-stack. (L) Autocorrelation function of the periodic region shown in K. (M) Lateral view of the tail of an *unc-70(FRT-flanked)* animal at the L4 stage after *unc-70* excision specifically in the epidermis, showing the localization of reconstituted epidermal SPC-1::GFP_x7_ within this tissue. (N) 3D-SIM image showing localization of reconstituted epidermal SPC-1::GFP_x7_, in a region surrounding the axon of the mechanosensory neuron ALM in an *unc-70(FRT-flanked)* animal at the L4 stage after *unc-70* excision specifically in the epidermis. Left panel is a lateral view, right panel an orthogonal perspective of the projected z-stack. (O) Autocorrelation function of the periodic region shown in D.

To determine if this periodic epidermal spectrin scaffold surrounding axons was required for the maintenance of axonal integrity, we sought to specifically disrupt epidermal *unc-70/ß-Spectrin*. To achieve this, we engineered a conditional allele of *unc-70/ß-Spectrin* flanked by FLP recognition target (FRT) sites using CRISPR-Cas9 (Fig. 3i). Importantly, when *unc-70/ß-Spectrin* was excised specifically in the epidermis, SPC-1::GFP_x7_ no longer localized to the epidermal membrane adjacent to motor and sensory axons (Fig. 3j & m). Following epidermal excision of *unc-70/ß-Spectrin* we observed no periodicity of SPC-1::GFP_x7_ surrounding axons (Fig. 3k-l & n-o). We then tested the effect of tissue-specific FLP-mediated excision of this region by driving FLP expression under the control of cell-specific promoters for the epidermis (SKIN::FLP = *Pdpy-7::FLP*), body wall muscle (MUSCLE::FLP = *Pmyo-3::FLP*), and nervous system (NEURONAL::FLP = *Prgef-1::FLP*) and visualized the morphology of mechanosensory and motorneurons. Epidermal excision of *unc-70/ß-Spectrin* caused axonal breaks in the PLM mechanosensory neurons as well as GABAergic motorneurons (Fig. 4a-h) and resulted in an uncoordinated body movement defect (Fig S3a); in contrast, pan-neuronal excision did not cause significant axonal damage (Fig. 4a-h), but did cause an uncoordinated body movement defect (Fig S3a). Similarly, excision of *unc-70/ß-Spectrin* cell-specifically in mechanosensory neurons (PLM::FLP = *Pmec-7::FLP*), GABAergic motorneurons (GABA::FLP = *Punc-47::FLP*) or body wall muscle (MUSCLE::FLP = *Pmyo3::FLP*) did not cause axonal damage (Fig. 4a-b). Despite observing almost no axonal damage in animals with neuronal excision of *unc-70/ß-Spectrin*, we did observe some axonal thinning and buckling in the axons of mechanosensory neurons, consistent with defects in axonal tension and mechanical strain resistance that have been observed previously (Fig. 4e) ^3,7^. Consistent with these axonal defects being the result of epidermal deletion of *unc-70/ß-Spectrin*, epidermal expression of wild-type UNC-70/ß-Spectrin was able to rescue motorneuron defects (Fig S3b) and axonal breaks in PLM mechanosensory neurons (Fig S3c). Pan-neuronal expression of wild-type UNC-70/ß-Spectrin provided a mild reduction to motorneuron defects (Fig S3b), whereas expression in mechanosensory neurons did not alter the penetrance of axonal breaks in PLM (Fig S3c). We observed a similar epidermal-specific effect of conditional deletion of *spc-1/α-Spectrin* on axonal integrity in motorneurons and the PLM mechanosensory neuron (Fig S3d-f). Together these results support the notion that the epidermal spectrin scaffold is necessary to protect axons from damage.

**Fig. 4.**
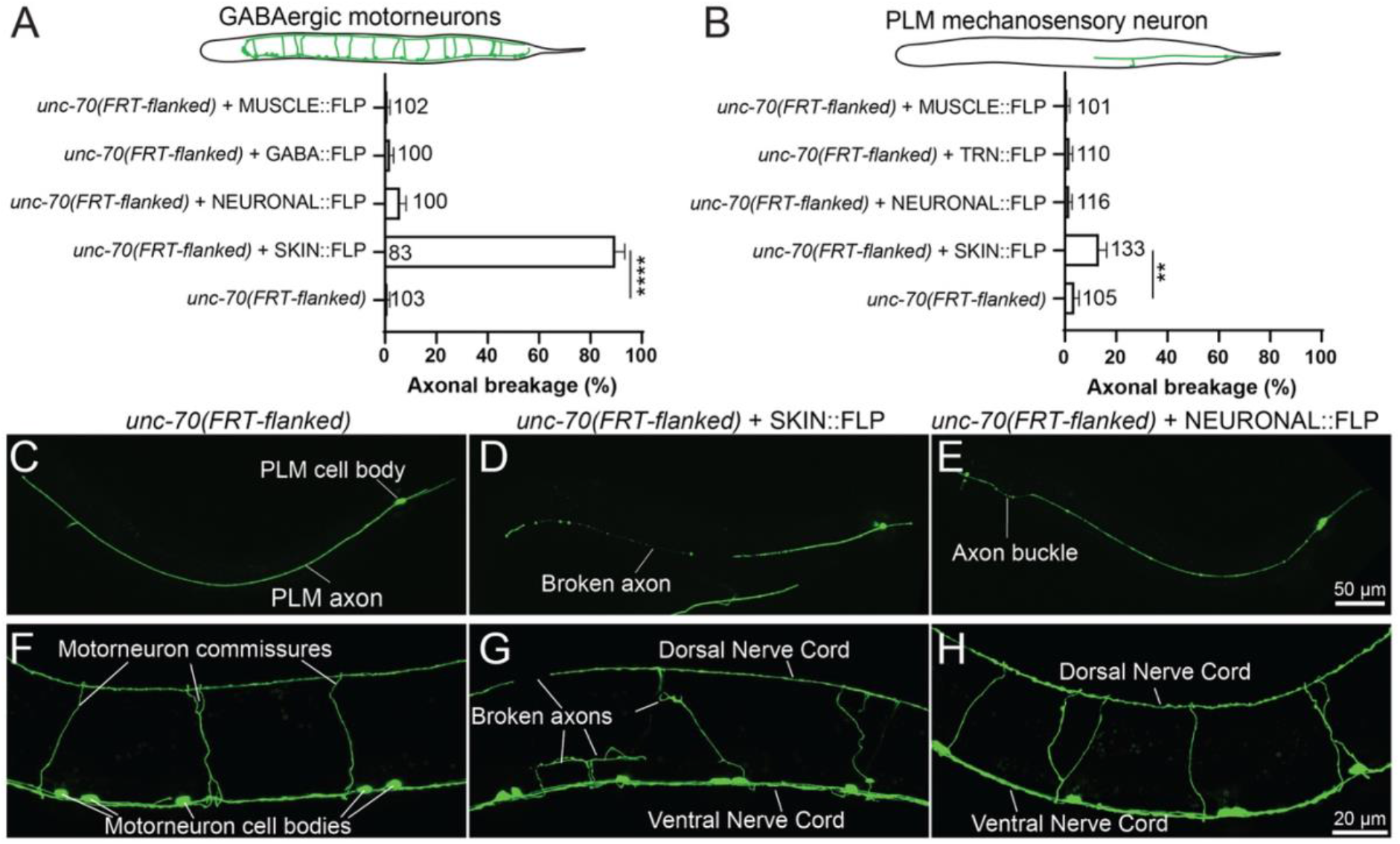
UNC-70/ß-Spectrin functions non-cell autonomously in the epidermis to maintain axonal integrity. (A) Quantification of axonal breakage in the commissures of DD and VD GABAergic motorneurons following tissue-specific excision of *unc-70/ß-Spectrin* in various tissues (Muscle = body wall muscles; GABA = GABAergic neurons; Neuronal = pan-neuronal; Skin = epidermis). (B) Quantification of axonal breakage in the PLM mechanosensory neurons following tissue-specific excision of *unc-70/ß-Spectrin* in various tissues (TRN = touch receptor neurons). (C-E) Images showing the morphology of the PLM neurons in control *unc-70(FRT-flanked)* animals (C), as well as after tissue-specific excision in the epidermis (D) and in the entire nervous system (E). (F-H) Images showing the morphology of DD and VD GABAergic motorneuron commissures in control *unc-70(FRT-flanked)* animals (F), as well as after tissue-specific excision in the epidermis and in the entire nervous system (H). **** in B indicates a P-value of < 0.0001 and ** in C indicates a P-value of 0.0032. P-values were determined from a one-way ANOVA with a Tukey multiple-comparison of proportions. Data in A and B are presented as mean ± SEM of the penetrance calculated using a uniform prior probability distribution and the number of animals scored is indicated on graphs.

In summary, our results support a model in which epidermal spectrin forms a periodic scaffold, which we refer to as the “epidermal MPS”, surrounding the nervous system where it is required to maintain axonal integrity. The unique structure of the axonal MPS has fortified the pervasive model that spectrin protects axons from damage by functioning intrinsically within the axon to resist mechanical strain. We observed a periodic localization of spectrins within the epidermis in regions adjacent to axons with a periodicity similar to that of the axonal MPS. This organization has previously been observed to be preferentially formed in axons compared to dendrites and rarely seen in localized regions of glial-cell processes ^1^. The majority of the epidermis of *C. elegans* is a syncytium containing 139 nuclei that envelops the whole body, except for the head and tail ^12^. The restricted periodic localization we observed corresponds precisely to the location of adjacent and enveloped axons, outlining the location of the nervous system within this tissue. Given the proposed roles for the spectrin cytoskeleton in cell signaling, cell adhesion and mechanical resistance, the epidermal MPS could function in multiple cellular processes to regulate the integrity, function, and development of adjacent axons.

## Materials and Methods

### Strains

All nematodes were cultured on nematode growth medium (NGM) plates seeded with *Escherichia coli* OP50 according to standard methods ^13^. Experiments were performed on L4 stage, or 1-day-old adults, raised at 20 °C. 1-day-old adults were selected 24 h prior to visualization at the L4 stage. The wild-type N2 Bristol strain was used with the alleles and transgenes listed in Table S1.

### Molecular biology

Standard molecular biology techniques were used ^14^. The *Pdpy-7::spGFP*_*1-10*_ plasmid was generated by excising an spGFP_1-10_ fragment from a *Pmec-4::spGFP*_*1-10*_ plasmid using BamHI and KpnI restriction enzymes, and cloning it downstream of the *Pdpy-7* promoter in a *Pdpy-7::unc-54-3’ UTR* plasmid. The *Prgef-1::unc-70cDNA* plasmid was generated by excising an *unc-70cDNA* fragment from a *Pdpy-7::unc-70cDNA* plasmid using SmaI and NheI restriction enzymes, and cloning it downstream of the *Prgef-1*promoter in a *Prgef-1::unc-54-3’ UTR* plasmid.

### CRIPSR-Cas9 gene editing

FRT insertions and spGFP_11×3_ knock-in animals were generated by microinjection of CRISPR–Cas9 protein complexes. Injection mixes contained 1.525 μM ALT-R Cas9 (Integrated DNA Technologies), 1.525 μM tracRNA (Integrated DNA Technologies), and 1.525 μM ALT-R crRNA (Integrated DNA Technologies) with 6 μM single-stranded DNA (FRT insertions; Integrated DNA Technologies) or 0.3-0.5 μM of PCR-amplified repair templates (spGFP_11×3_). All repair templates were designed with 75 base pair homology to flanking genomic regions. PCR-repair templates were amplified using ultramer oligonucleotides. Sequences of all crRNAs and oligonucleotides used are listed in Tables S2 and S3, respectively. All genomic edits were verified by PCR and Sanger sequencing. For spGFP_11×3_::UNC-70 knock-in animals, a repair template carrying spGFP_11×7_ was used, but the resultant genomic edit contained only x3 tandem repeats of spGFP_11_.

### Phenotypic analysis

Axonal breakage in PLM neurons was scored in 1-day-old adult animals containing the *zdIs5(Pmec-4::GFP)* transgenes and selected 24 h prior to visualization at the L4 stage. Axonal breakage in motorneuron commissures was scored at the L4 stage in animals containing the *oxIs12(Punc-47::GFP)* transgene. All raw data is included in Table S4.

### Microscopy and analysis

Animals were visualized by mounting them on 4% agar pads after immobilization with 0.05% tetramisole hydrochloride. Phenotypic analysis was performed on an upright Zeiss AxioImager A1 microscope (Carl Zeiss AG). Confocal imaging was performed at the Queensland Brain Institute’s Advanced Microscopy Facility using a spinning disk confocal microscope (Marianas, Intelligent Imaging Innovations), equipped with a confocal scanner unit (CSU-W1, Yokogawa Electric Co.) built around an Axio Observer body (Z1, Carl Zeiss AG) and fitted with an sCMOS camera (ORCA-Flash4.0 V2, Hamamatsu Photonics) and SlideBook v6.0 software (3i) using a 100×/1.46 NA oil-immersion objective with sampling intervals x,y = 63 nm and z = 130 nm. Images were deconvolved with Huygens Professional v18.04 run on a GPU-accelerated computer (3× NVIDIA® Tesla® V100) using the CMLE algorithm, with a signal to noise ratio of 20, background of 100, and 40 iterations. Images were exported as 16-bit TIFs and further processed in Fiji v1.52p ^15^ to obtain YZ-plane orthogonal views. Microinjections were performed using standard methods ^16^, with an inverted Zeiss AxioObserver microscope equipped with differential interference contrast, a Narishige needle holder, and a pressurized air pump. Three-dimensional structured illumination microscopy (3D-SIM) was performed on a Zeiss Elyra PS.1 SIM/PALM/STORM system generously supported by the Australian Government through an ARC LIEF grant (LE130100078) using an Alpha Plan-Apochromat 100×/1.46 NA oil-immersion objective, and SIM processing was performed using ZEN Black 2012 SP5 version 14.0.0.0 (Carl Zeiss AG). 10 frame z-stacks were acquired with a z-interval of 101 nm, and SIM processing performed. GFP fluorescence was visualized with a PCO Edge sCMOS camera (PCO AG) and a 488 nm laser (15 % power, 300 ms exposure time, 0.41 mm grating, 3 rotations). mCherry was visualized with a 561 nm laser (10 % power, 100 ms exposure time, 0.52 mm grating, 3 rotations). SIM processing was completed with a noise filter value ranging from - 4.2373 to -5.1796 and sectioning of 100/83/83. Autocorrelation of periodic spectrin localization was performed using Fiji v1.52p ^15^ and Matlab (Mathworks).

## Acknowledgments

We thank R. Amor and A. Gaudin for support with microscopy; K. Wiesshart for technical assistant with 3D-SIM; P. Bartlett, P. Bazzicalupo, J. Götz, and R. Tweedale, for comments on the manuscript; colleagues in the Hilliard laboratory for insightful discussions and comments. Some strains were provided by the CGC, which is funded by NIH Office of Research Infrastructure Programs (P40 OD010440), and the International *C. elegans* Gene Knockout Consortium.

## Author contributions

SC and IBP performed experiments, collected, and analyzed data. SC, IBP, and MAH conceptualized and designed experiments, discussed results, interpreted data, and wrote the manuscript.

## Data availability

All data are available in the main text or the Supplementary material.

## Supplementary Material

**Fig. S1.**
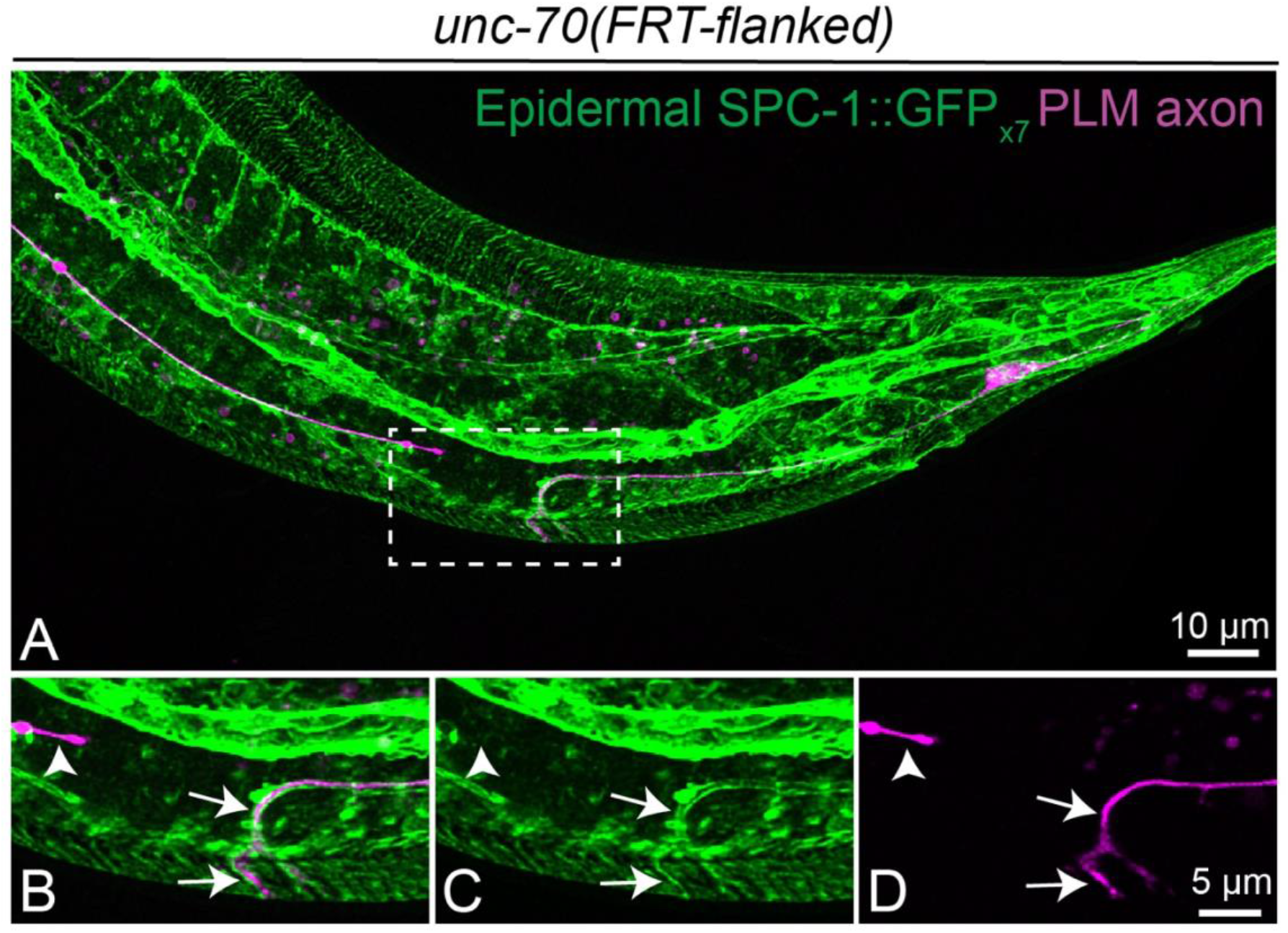
Epidermal SPC-1/α-Spectrin localizes to regrowing axons after injury. (A) Lateral view of the tail of an *unc-70(FRT-flanked)* animal, 48 hours after axotomy, showing the localization of reconstituted epidermal SPC-1::GFP_x7_ (SPC-1::spGFP_11×7_ + SKIN::spGFP_1-10_) with a red fluorescent marker labeling the PLM mechanosensory neuron. (B-D) High magnification view of the boxed region in A. Arrows indicate epidermal SPC-1::GFP_x7_ localization around the proximal regrowing fragment of the PLM axon. Arrowheads indicate a region where SPC-1::GFP_x7_ localization is absent from the separated distal axonal fragment of the PLM neuron.

**Fig. S2.**
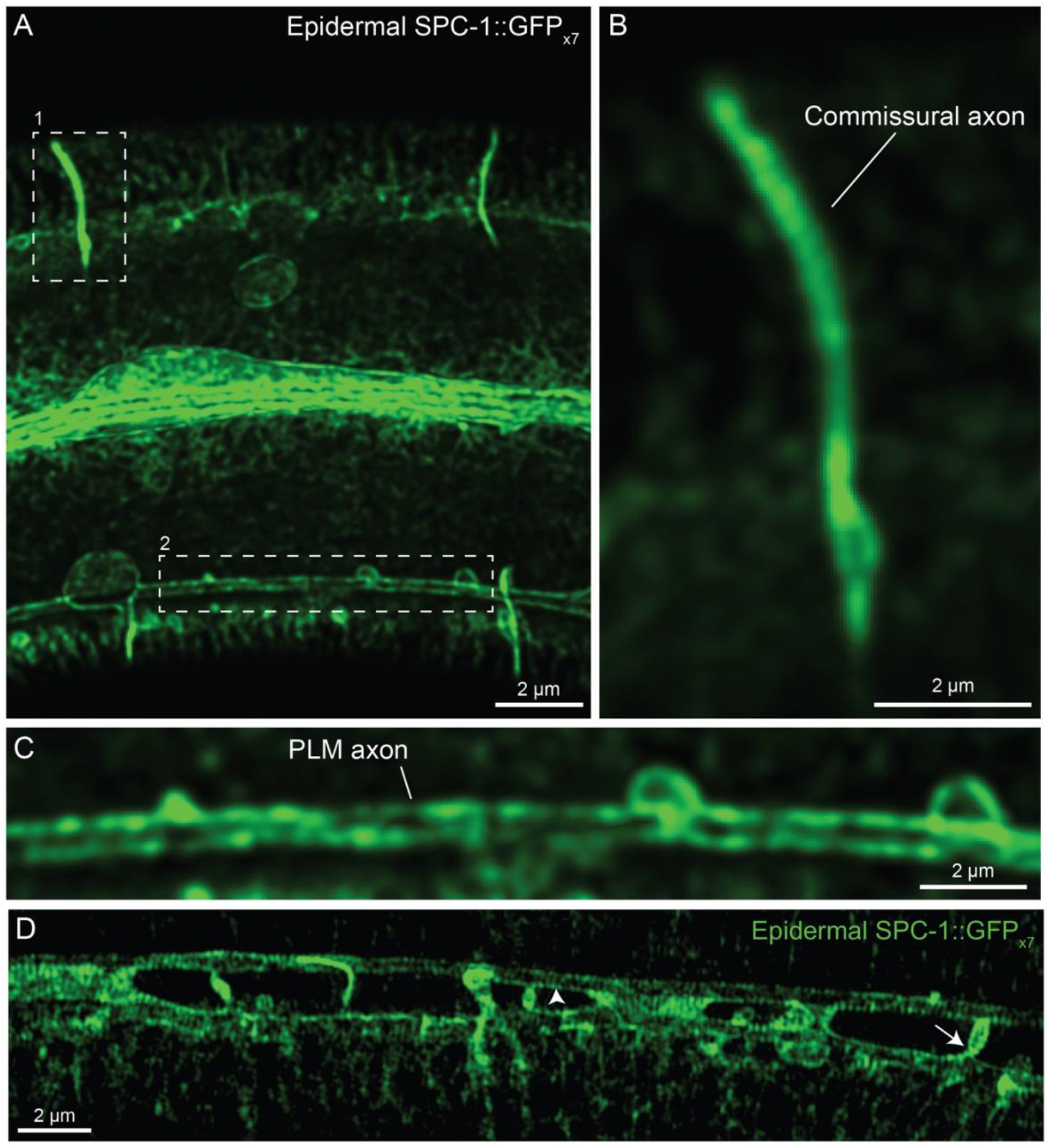
Epidermal SPC-1/α-Spectrin localizes to discontinuous periodic structures surrounding axons. (A) Lateral view of a wild-type animal at the L4 stage showing the localization of reconstituted epidermal SPC-1::GFP_x7_ (SPC-1::spGFP_11×7_ + SKIN::spGFP_1-10_). (B) High-magnification view of boxed area 1 in A showing a region adjacent to a commissural axon. (C) High-magnification view of boxed area 2 in A showing a region surrounding the PLM mechanosensory neuron. (D) 3D-SIM image showing localization of reconstituted epidermal SPC-1::GFP_x7_ in a region surrounding the PLM axon. Arrowhead indicates periodic localization orthogonal to a horizontal structure, arrow indicates periodic structures orthogonal to a vertical structure.

**Fig. S3.**
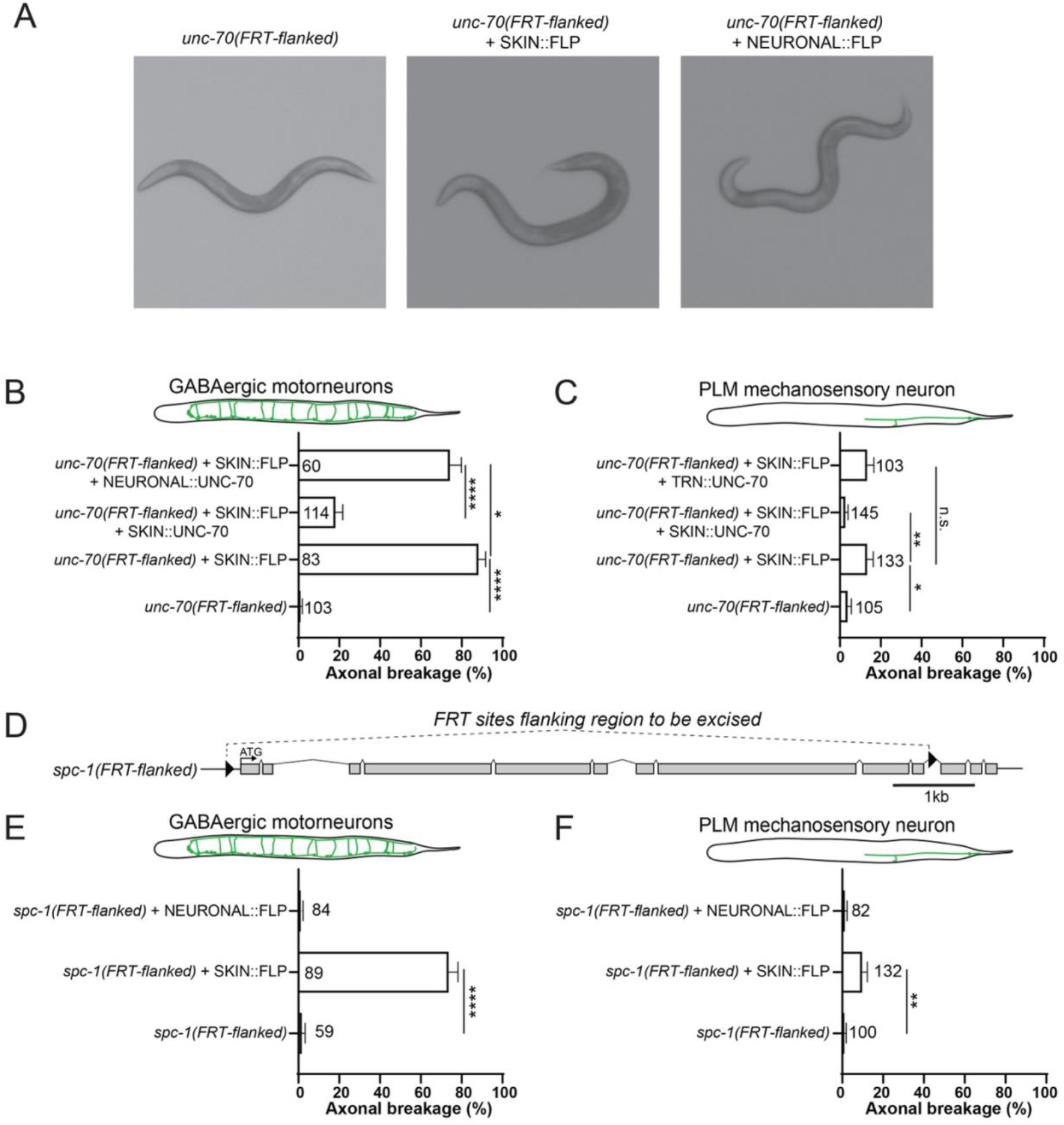
Epidermal excision of UNC-70/ß-Spectrin causes axonal breaks and movement defects. (A) Images showing the body posture of control animals, and animals with epidermal or pan-neuronal excision of *unc-70*. (B) Quantification of axonal breakage in the commissures of DD and VD GABAergic motorneurons, following tissue-specific excision of *unc-70/ß-Spectrin* from epidermal tissue with tissue specific rescue of wild-type UNC-70 from either neurons or epidermis (Neuronal = pan-neuronal; Skin = epidermis). (C) Quantification of axonal breakage in the PLM mechanosensory neurons, following tissue-specific excision of *unc-70/ß-Spectrin* from epidermal tissue with tissue specific rescue of wild-type UNC-70 from either mechanosensory neurons or epidermis (TRN = touch receptor neurons; Skin = epidermis). (D) A schematic depicting a conditional *spc-1/a-Spectrin* allele containing flippase recognition target (FRT) sites for tissue-specific flippase-dependent excision. (E) Quantification of axonal breakage in the commissures of DD and VD GABAergic motorneurons following tissue-specific excision of *spc-1/a-Spectrin* from either neurons or epidermis. (F) Quantification of axonal breakage in the PLM mechanosensory neurons following tissue-specific excision of *spc-1/a-Spectrin* from either neurons or epidermis. Statistical differences in B-C & E-F were determined using one-way ANOVA with a Tukey multiple-comparison. Data are presented as mean ± SEM of the penetrance calculated using a uniform prior probability distribution and the number of animals scored is indicated on graphs.

**Table S1.**
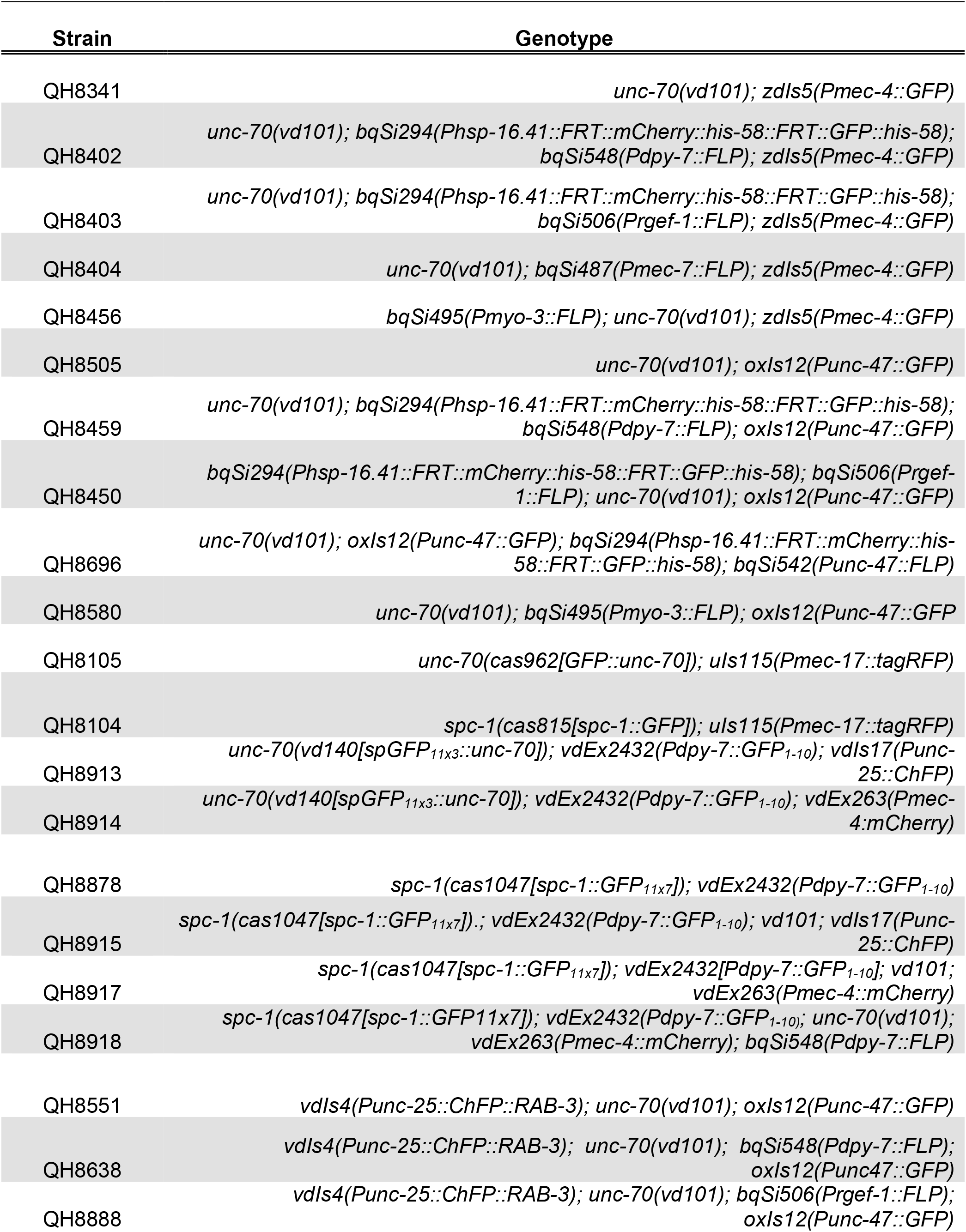
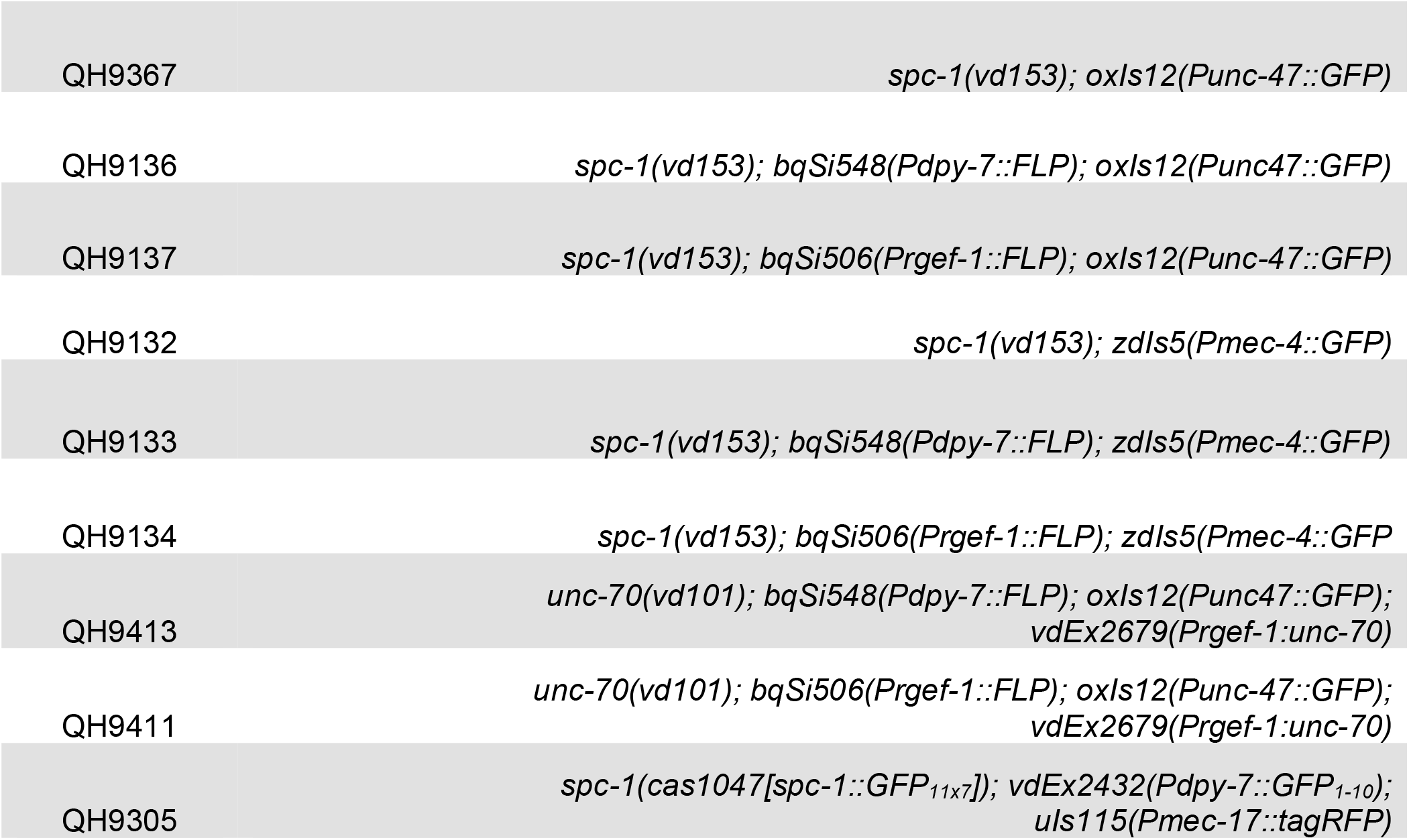
*C. elegans* strains used.

**Table S2.**
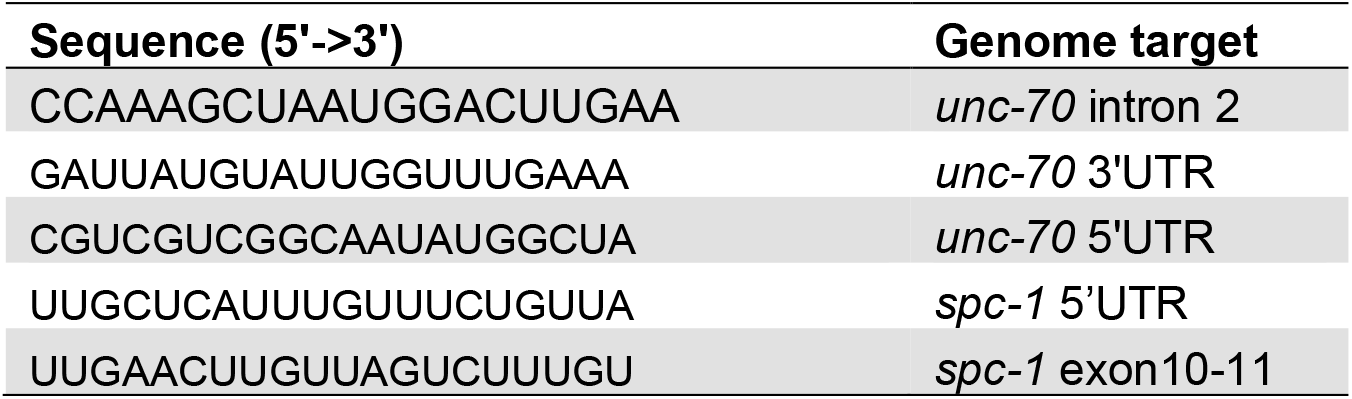
crRNA sequences used for CRISPR-Cas9 gene editing.

**Table S3.**
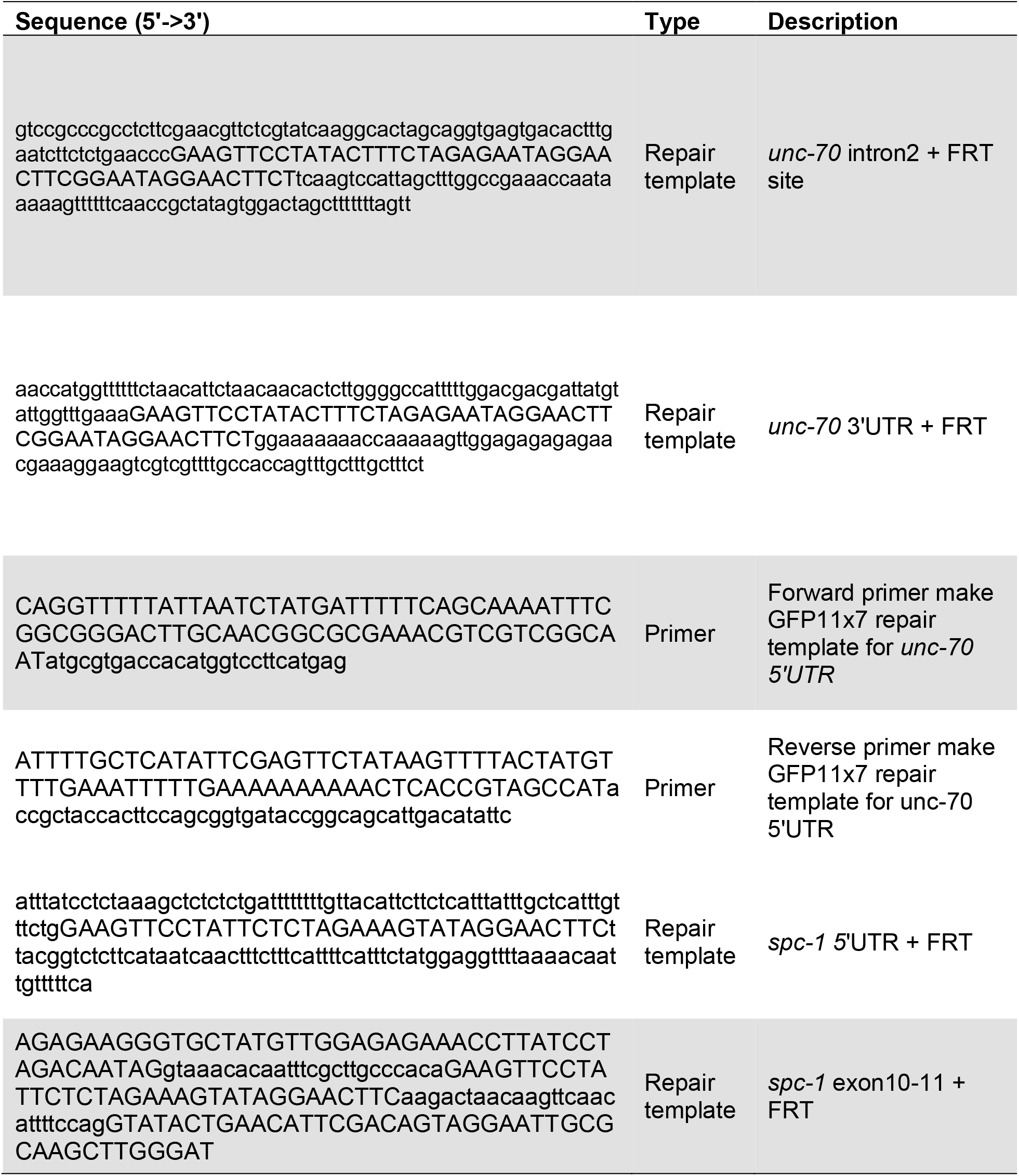
Ultramer oligonucleotides used for CRISPR-Cas9 gene editing.

**Table S4.** Raw data of axonal breakage phenotypes.

| Strain | Genotype | Stage | Commissure break | n |
| --- | --- | --- | --- | --- |
| QH8505 | *oxls12; unc-70(vd101)* | L4 | 0 | 103 |
| QH8459 | *oxls12; unc-70(vd101); bqSi548; bqSi249* | L4 | 60 | 66 |
| QH8450 | *oxls12; unc-70(vd101); bqSi506; bqSi249* | L4 | 5 | 100 |
| QH8696 | *oxls12; unc-70(vd101); bqSi452* | L4 | 1 | 100 |
| QH8580 | *oxls12; unc-70(vd101); bqSi495* | L4 | 0 | 102 |
| QH9367 | *oxls12; spc-1(vd153)* | L4 | 0 | 59 |
| QH9136 | *oxls12; spc-1(vd153); bqSi548* | L4 | 66 | 89 |
| QH9137 | *oxls12; spc-1(vd153); bqSi506* | L4 | 0 | 84 |

|  |  |  | Transgenic |  | Non- Transgenic |  |
| --- | --- | --- | --- | --- | --- | --- |
| Strain | Genotype | Stage | Commissure break | n | Commissure break | n |
| QH9413 | *oxls12; unc-70(vd101); bqSi548; vdEx2679(Prgef-1:unc-70)* | L4 | 45 | 60 | 35 | 40 |
| QH9411 | *oxls12; unc-70(vd101); bqSi506; vdEx2679(Prgef-1:unc-70)* | L4 | 1 | 43 | 1 | 31 |

| Strain | Genotype | Stage | PLM Break | n |
| --- | --- | --- | --- | --- |
| QH8341 | *zdls5; unc-70(vd101)* | 1DOA | 3 | 105 |
| QH8402 | *zdls5; bqSi548; unc-70(vd101);bqSi249* | 1DOA | 17 | 133 |
| QH8403 | *zdls5; unc-70(vd101); bqSi506* | 1DOA | 1 | 116 |
| QH8404 | *zdls5; bqSi487; unc-70(vd101)* | 1DOA | 1 | 110 |
| QH8456 | *zdls5; unc-70(vd101); bqSi495* | 1DOA | 0 | 101 |
| QH9173 | *zdls5; unc-70(vd101); vdSi3; bqSi548* | 1DOA | 2 | 145 |
| QH9104 | *zdls5; vd101; bqSi548; vdSi7* | 1DOA | 13 | 103 |
| QH9132 | *spc-1(vd153); zdls5* | 1DOA | 0 | 100 |
| QH9133 | *spc-1(vd153); bqSi548; zdls5* | 1DOA | 12 | 132 |
| QH9134 | *spc-1(vd153); bqSi506; zdls5* | 1DOA | 0 | 82 |

